# From Genomics Alterations to Expression Dynamics: A Hierarchical Multi-Omics Integration Framework with gINTomics

**DOI:** 10.1101/2025.10.07.680960

**Authors:** Angelo Velle, Francesco Patanè, Chiara Romualdi

## Abstract

Despite the abundance of multi-omics integration methods, few adopt a hierarchical approach, which is crucial for understanding how different genomic alterations influence downstream molecular processes. We present `gINTomics`, a hierarchical computational framework that captures directional biological information flow across genomic, transcriptomic, and epigenomic layers. Unlike existing meta-dimensional approaches, `gINTomics` follows a multi-staged paradigm reflecting biological hierarchy, modeling molecular modifications by linking genomic alterations to downstream transcriptional consequences. The framework offers flexible integration strategies for different omics combinations and tailored statistical modeling with optional Random Forest for variable selection. A key feature is the comprehensive interactive Shiny application that provides intuitive visualization through interactive plots and circos diagrams, enabling users to explore complex regulatory relationships across multiple analytical perspectives. Validation on TCGA ovarian cancer data confirmed the framework’s effectiveness in identifying biologically relevant regulatory mechanisms and prognostic biomarkers. Available as a Bioconductor package, gINTomics provides researchers with a powerful and accessible tool for mechanistic multi-omics integration and biomarker discovery.

## BACKGROUND

The advent of high-throughput technologies for omics data acquisition has led to an unprecedented accumulation of heterogeneous biological data, encompassing gene expression, DNA methylation, copy number variations, and more. These large-scale, multi-modal, high-resolution datasets offer a multidimensional view of cellular systems and hold great potential for unraveling the molecular mechanisms underlying complex phenotypes, including cancer^1–5^. To harness this potential, the scientific community has developed a wide array of computational methods aimed at integrating multi-omics data. While most existing integration strategies focus on tasks such as patient stratification or cancer subtype discovery, fewer methods are explicitly designed to explore mechanistic relationships—e.g., linking genomic alterations to downstream transcriptional consequences—or to identify interpretable biomarkers through biologically informed models.

Over the past decade, a substantial body of literature has emerged in the field of multi-omics integration^6–9^, leading to various methodological frameworks that can be divided into general and broad classification schemes^6–9^. Among the earliest and most prominent are: multi-staged (or hierarchical) integration and meta-dimensional analysis^9^.

Meta-dimensional approaches (which can be further subdivided into finer subclasses) aim to integrate all omics layers simultaneously, treating them as equally informative features in a joint multivariate model—often without incorporating biological priors. Within this category, several R packages have been developed, particularly in the context of cancer subtyping (see Supplementary Table 1 for an overview of available methods^10–25^).

For example, MOFA^10^ is a computational approach that identifies the main sources of variation in multi-omics datasets. It uses latent factor analysis to extract hidden factors representing biological and technical sources of variability, and distinguishes between dimensions of heterogeneity that are shared across data modalities and those specific to individual modalities. Conceptually, MOFA can be seen as a generalization of principal component analysis for multi-omics data. The extracted factors enable downstream analyses such as subgroup discovery, data imputation, and outlier detection.

Mixomics^11^ is another R package dedicated to the multivariate analysis of biological datasets, with a particular focus on data exploration, dimensionality reduction, and visualization. It offers several statistical methods that integrate multiple datasets simultaneously to explore associations across different omics layers. Some of its latest approaches are based on Projection to Latent Structure (PLS) models, used for classification, multi-omics integration across datasets, and molecular marker detection.

MOVICS^12^ provides a unified interface for ten multi-omics integrative clustering algorithms, and includes frequently used downstream analyses in cancer subtyping—such as characterizing and comparing discovered subtypes and validating them in external cohorts. Lastly, MOSClip^14^ performs multi-omics integration at the pathway level by applying omic-specific dimensionality reduction techniques, followed by multivariate survival modeling to identify pathways associated with survival outcomes.

In contrast, the multi-staged approach breaks the integration process into a sequence of analytical steps, typically reflecting the presumed biological hierarchy among omics layers— for instance, treating genomic alterations as upstream drivers of transcriptional or epigenetic changes. Within this category, only a few approaches exist, mainly based on integrative Bayesian frameworks^25^ or regression models^23,24^. In some cases, the corresponding software is no longer available or was never implemented in a publicly accessible platform. In others (e.g., COSMOS), the genomic layer is used solely to filter for reliable transcriptional relationships (e.g., TF–target interactions), rather than to interpret observed expression changes in terms of upstream molecular alterations.

Beyond the computational strategy adopted, another major challenge in multi-omics integration lies in summarizing and visualizing the results. The sheer volume of inferred molecular relationships and phenotype associations demands dynamic and efficient visualization tools capable of unlocking the full potential of the information generated by these approaches.

In this work, we propose `gINTomics`, a novel hierarchical multi-omics integration framework that builds on the multi-staged paradigm. Our method is specifically designed to capture the directional flow of biological information across omics layers, enabling interpretable modeling of molecular cascades and more accurate prediction of complex phenotypes.

To demonstrate its ability to detect the impact of genomic and transcriptional alterations on gene expression, we apply `gINTomics` to two case studies derived from publicly available multi-omics data from the TCGA Ovarian Cancer cohort.

## MATERIALS AND METHODS

### Rationale and Structure of the package

`gINTomics` is a hierarchical computational framework designed for the analysis of multi-omics datasets. It enables users to explore data from multiple perspectives, offering an intuitive and user-friendly interface to delve into results. The most distinctive feature of `gINTomics` is its ability to identify associations between the expression of a target gene and its regulators, while accounting for the influence of genomic alterations—such as Copy Number Variations (CNVs) and DNA methylation—on those regulators. To support this goal, `gINTomics` allows for both two-omics and multi-omics data integration (Figure 1).

**Figure 1.**
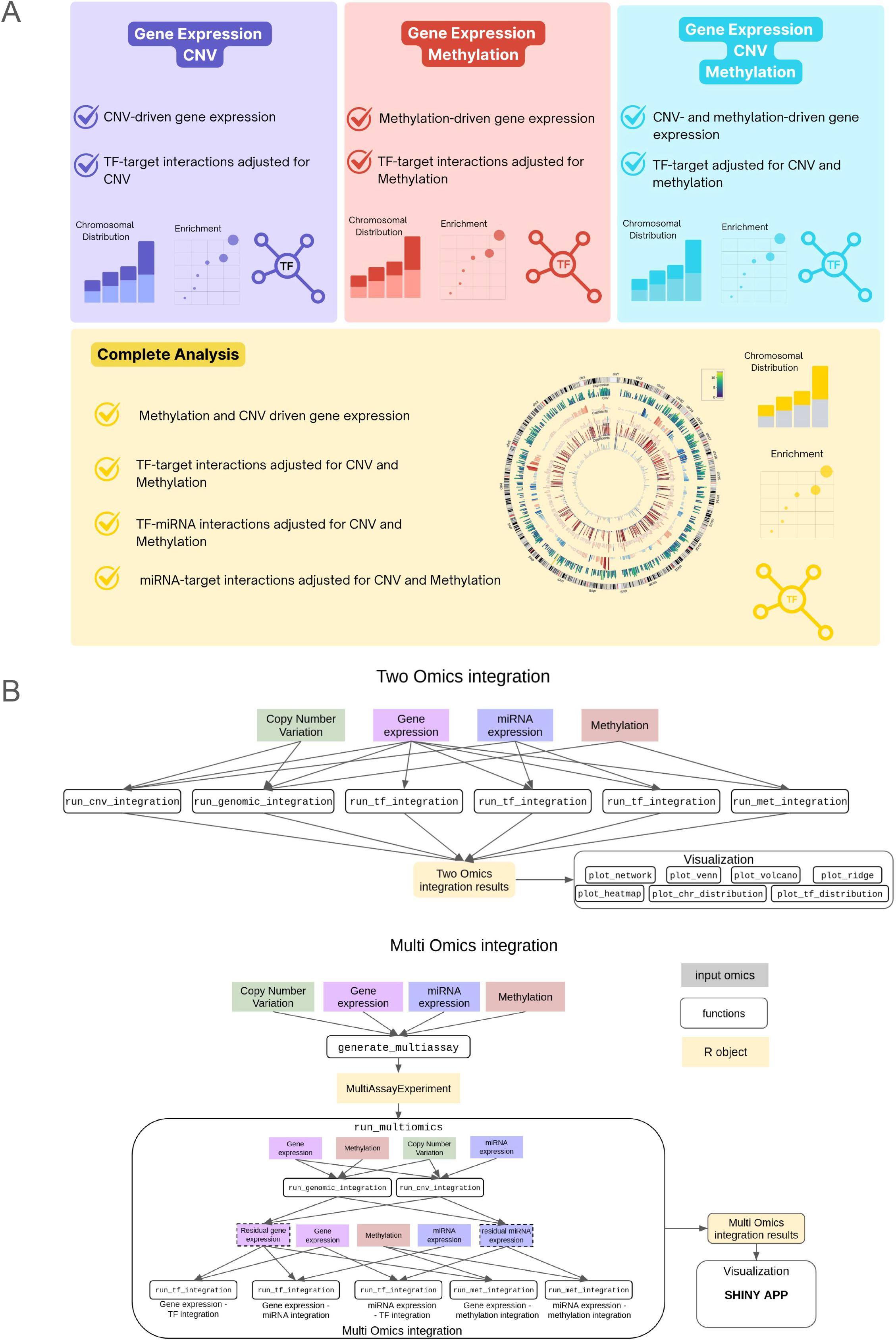
gINTomic structure overview. **(A)** Comprehensive representation of integration approaches available for various omics data combinations. **(B)** Schematic workflow illustrating available functions for data integration and visualization, including their respective input data requirements.

The workflow for all possible integrations is also illustrated in Figure 1A. In each integration panel, a statistical model is applied where the response variable is always gene/miRNA expression levels, while the regulators (e.g., transcription factors expression, CNV status, methylation status) are included as covariates. All models are gene-specific, meaning each gene/miRNA has its own model with its corresponding covariates.

Different functions are available for two-, or multi-omics integration, depending on the data provided by the user (see Figure 1B and Table 1).

**Table 1.** gINTomic functions. A list of `gINTomic` functions applicable to the omics data types, highlighting the available approaches for multi-omics integration.

Integration functions exploit different statistical models depending on the input data type: the negative binomial regression (as implemented in the `edgeR` package^26^) and linear regression (as implemented in the `stats` package^27^). Moreover, to handle situations in which the number of regulators is too high, `gINTomics` supports a Random Forest variable selection that will keep just the most important covariates defined by the mean decrease in accuracy with an Out-Of-Bag method of the `randomForest` R package^28^ (after the Random Forest selection the number of covariates will be the 40% of the number of samples).

A multi-class design is implemented where the user can provide labels for the separation of samples into multiple classes. In this case `gINTomics` will analyse the data for all classes together, highlighting the results of the models involving differentially expressed genes.

In order to make the results more interpretable, `gINTomics` provides many graphical representations of the results as well as a powerful shiny app that allows users to navigate interactively into models’ results. The shiny app can be started with the *run_shiny* function.

### Implementation

For two omics integration, the input data should be provided as matrices or data frames. For multi omics integration, the input data should be a *MultiAssayExperiment*^29^ object. To simplify the creation of this object, the package includes a dedicated function called *create_multiass*ay which accepts a list of matrices or *SummarizedExperiment*^30^ objects — one for each omic type (with a *minimum of two*). This function automatically constructs a *MultiAssayExperiment* object that consolidates all input data and is ready to be used in the integration model. Expression data can be either raw or normalized; if raw data are provided, the package will handle normalization internally. The integration functions include an argument that allows users to specify the normalization method and whether it should be applied.

#### Integration Functions

*run_cnv_integration:* integration of gene/miRNA expression with Copy Number Variations (CNV). The goal of this function is to assess the impact of CNV alterations on gene or miRNA expression levels. It requires as input two matrices or data frames: the first contains expression values, where rows represent samples and columns correspond to response variables (e.g., genes or miRNAs); the second contains CNV data, with rows again representing samples and columns representing the covariates of interest. Users must specify whether the expression data originate from sequencing experiments—in which case a negative binomial regression model will be applied. Otherwise, a linear regression model is used. All integration models support parallel execution via the BPPARAM argument; by default, models run on a single core. For each gene, the model estimates the regression coefficient associated with the CNV covariate and its corresponding p-value. A positive coefficient indicates a concordant association between CNV and expression (i.e., higher CNV values are associated with increased expression), while a negative coefficient indicates an inverse relationship.

*run_met_integration:* This function is analogous to the one used for CNV integration but is specifically designed to model the relationship between gene expression and DNA methylation. Its primary goal is to identify the impact of methylation alterations on changes in gene expression.

*run_genomic_integration:* integration of gene expression with both CNV and methylation together. The goal of this function is to identify the effect of both CNV and methylation alterations on expression modifications. The function has the same structure of the previous ones and takes as input three matrices, one for gene expression, one for CNV data and one for methylation data. For each gene the model will estimate the value of the parameters for CNV and for methylation along with their p values.

*run_tf_integration:* transcriptional integration suitable for the TF-target gene, TF-miRNA and miRNA-target integrations. The aim of the function is to identify the combined TF activity on their target gene expression. The integration between target genes and their Transcription Factors (TF) can be performed by setting as “tf” the type argument. The function takes as input two matrices or data frames: the first one is the expression values for each target gene, rows represent samples, while each column represents the different response variables of the models; the second one is the expression of the TFs, rows represent samples, while columns represent the different covariates. If the first matrix contains both target genes and TFs, the second matrix is not needed. The function will automatically download the interactions between target genes and TFs from the `OmnipathR` R package^31^, unless they are provided by the user through the *interactions* argument.

The *run_tf_integration* function can integrate target miRNA expression data with transcription factor (TF) expression data by setting the *type* argument to “*tf_miRNA*”. It requires two input matrices or data frames: the first should contain expression values for each target miRNA, with rows as samples and columns as response variables; the second should contain expression values for each TF, with rows as samples and columns as covariates. By default, the function uses a built-in interaction table from the TransmiR database^32^, unless the user provides a custom interaction list via the interactions argument. The species of interest must be also specified. Only literature curated interactions are considered.

Another integration mode supported by *run_tf_integration* analyzes the correlation between miRNA expression level and those of their target genes. This can be done by setting *type* equal to “miRNA_target” and operates similarly to the “tf” and “tf_miRNA” modes. The first input matrix should contain the expression values for each target gene, while the second one should contain the expression values for each miRNA. The function automatically retrieves interaction data via the `OmnipathR` R package unless the user supplies their own through the interactions argument.

For all these integration approaches, run_tf_integration estimates a coefficient and corresponding p-value for each regulator-target pair. The interpretation of coefficients aligns with other models: a positive coefficient indicates a concordant relationship, while a negative coefficient indicates an inverse association.

*run_multiomics:* This function performs a comprehensive integration by accepting a *MultiAssayExperiment* object as input and automatically detecting the available omics layers. Its goal is to identify the combined regulatory activity of transcription factors (TFs) on target gene expression while accounting for modifications due to CNV and DNA methylation. The workflow proceeds as follows: if both gene expression and genomic data (copy number variations and methylation) are available, they are first integrated using the *run_genomic_integration* function. The residuals from this model —representing gene expression values adjusted for CNV and methylation effects —are then used in subsequent analysis steps, as illustrated in Figure 1B. The integration models applicable to each data type are summarized in Table 1. The function returns an object of class *MultiOmics*, which is a list containing the results from each individual integration step.

#### Class Comparison

`gINTomics` also supports subsetting samples by class through the *classes* argument of the *run_multiomics* function. When this option is used, `gINTomics` performs a full analysis on all samples, highlighting the results for differentially expressed genes (DEGs) across the defined classes. The Class Comparison section of the Shiny app provides an intuitive interface to explore these results. Users can dynamically switch between available contrasts to browse the findings involving genes that are differentially expressed between classes, highlighting potential regulatory rewiring driven by class-specific differences. In addition, the Class Comparison section enables differential methylation and CNV analyses between any two selected classes (see the Shiny App and Visualization sections for details).

#### Shiny App and visualization

To facilitate model interpretation, `gINTomics` offers a comprehensive and interactive Shiny app that enables users to explore the results of all integration models in an intuitive and user-friendly way. The application can be launched using the *run_shiny* function, which takes as input the results of a multi-omics integration analysis.

The `gINTomics` visualizer is divided into four sections:

1. <u>Genomic Integration</u> for the results regarding copy number variations and methylation;
2. <u>Transcription Integration</u> for those regarding transcriptional networks (Transcription Factors and miRNA);
3. <u>Class Comparison</u> to highlight the results only for genes that are differentially expressed/altered among classes defined by the user;
4. <u>Complete Integration</u> for a comprehensive table with all the available results and a visual representation with a Circos plot for all the results;

The <u>Genomic Integration</u> section shows the estimates of significant parameters through ridgeline plots, volcano plots, and Venn diagrams, highlighting genes whose expression is significantly influenced by CNVs and/or methylation, with the possibility of showing their genomic location.

The <u>Transcription Integration</u> section reports the estimates of the significant parameters, focusing on genes whose expression is significantly affected by transcription factors (TFs). Additionally, we display significant TF–target gene associations using network plots, where TFs are connected to their targets via edges whenever a significant expression-based association is detected—taking into account genomic modifications if this information is provided by the user.

Both sections also provide an enrichment analysis to give a functional and biological interpretation of the results. The enrichment is performed with an Over-Representation analysis using the `clusterProfiler` R package^33^. In the Genomic sections we use the list of genes significantly regulated by CNV or methylation while in the Transcriptional section, we select the TF with the highest number of significant targets and we use them to run the enrichment. Many other plots are provided to investigate the relationship between variables, such as heatmaps, scatterplots and networks.

The <u>Class Comparison</u> section allows users to visualize and download the full list of differentially expressed genes for each contrast, and to perform Differential Methylation and CNV analyses. The Differential Methylation analysis accounts for the effect of CNVs (if available) on gene expression and uses the `MethylMix` R package^34^ to identify and visualize differentially methylated genes that significantly influence expression. The Differential CNV analysis is performed with a t-test on CNV values for each gene between the selected classes, providing a scatterplot to explore the correlation with gene expression and enabling result export in tabular format.

The <u>Complete Integration</u> section provides a circos plot that combines gene expression, CNV and methylation data (if available) to better visualize the integration. The section also contains a downloadable table with the results of all the integration performed by the package.

All the results summarized in the Shiny App are shown as figures or tables which are easily downloadable directly from the Shiny App. Finally, most of the plot available in the shiny app can be generated also outside the shiny environment, exploiting our plotting functions.

### Code and Software availability

`gINTomics` is available as a Bioconductor package (10.18129/B9.bioc.gINTomics)

### Case studies data

Ovarian Cancer data from the TCGA portal were used to demonstrate the potential of `gINTomics`, leveraging the extensive genomic alterations that characterize this tumor type. Specifically, two analyses were conducted: first, we analyzed the entire cohort of OV samples^35^ to explore the impact of CNV and methylation alterations on gene expression and biological processes. Only primary tumor data were analyzed, for a total of 385 samples. Second, we compared long-vs. short-term survival samples to investigate the rewiring of transcriptional networks in these two groups. Again, only primary tumors were considered. For the short-term survival group, we selected patients with a Progression Free Survival (PFS) shorter than 6 months (n=29), while for the long-term survival group, we selected samples with a PFS longer than 5 years (n=21).

Gene expression, Methylation, miRNA expression and clinical data were downloaded using the `TCGAbiolinks` R package^36^. GISTIC Copy Number Variations data for the same cohort were downloaded with the `curatedTCGAData` R package^37^. For gene expression, raw counts data were analyzed, exploiting the TMM normalization procedure provided by `gINTomics`. To assign a single methylation value to each gene, methylation beta values were clustered using the `MethylMix` R package. When more clusters were identified for a given gene, the highest value was assigned. For gene expression, 60660 genes were available, 24776 for CNV, 12404 for methylation, 1881 for miRNA expression and 1202 for miRNA CNV. Finally, the MultiAssay object was created exploiting the *create_multiassay* function of `gINTomics`.

## RESULTS and Discussion

Ovarian Cancer is one of the leading causes of cancer death among women. These tumors are characterized by widespread genomic alterations, which may impair the molecular characterization and make it very difficult to detect expression biomarkers^38,39^. Moreover only few of these genomic alterations are shared across patients and are likely to be key driving events, that’s why Ovarian Cancer represents a challenging test for our package to evaluate the impact of these alterations on gene expression and how these alterations might explain differences in clinical prognosis.

### gINTomics quantifies the impact of genomic and transcriptional alterations on gene expression in late stage ovarian cancer

Considering the entire cohort of OV samples (n=385), we found that 88% of the genes have the expression significantly regulated by CNV. Figure 2A illustrates the distribution of the p-values and of the coefficient estimates for CNV, highlighting that nearly all significant coefficients are positive. This indicates as expected higher expression levels for genomic gains and lower expression levels for losses. Using enrichment analyses we can evidence that, these genes are known to play a role in other cancers such as breast, gastric and skin cancer (as reported in Figure 2C) and seems to be involved in neuroactive ligand receptor interaction, cAMP signalling and ERB signalling, pathways known to influence cellular proliferation, angiogenesis and immune escaping.

**Figure 2.**
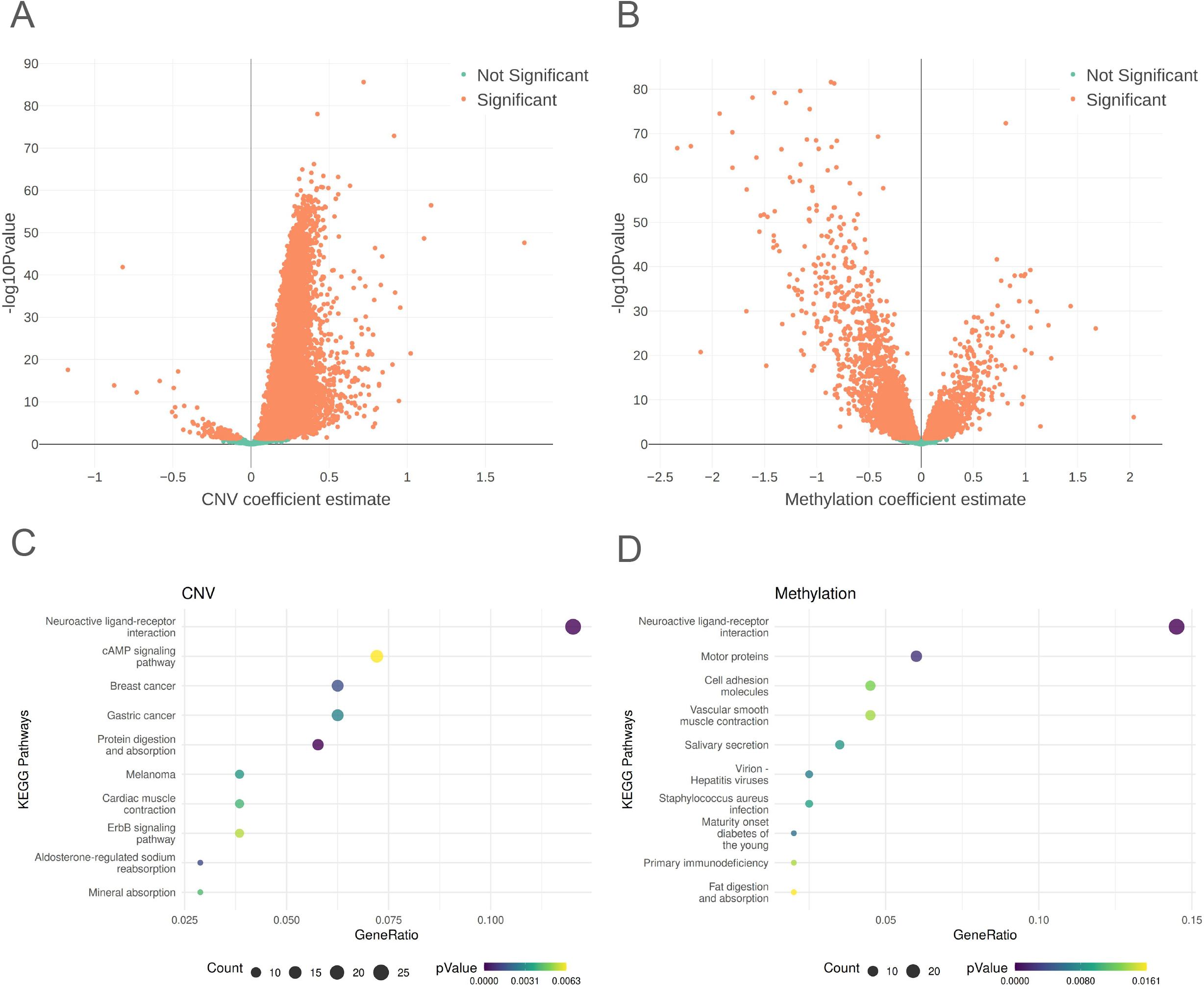
gINTomic analysis on the TGCA OV dataset. **(A)** Volcano plot of the coefficient estimates for copy number variations (CNVs) derived from the genomic integration model. The X-axis represents coefficient estimates, while the Y-axis shows the −log10 transformed P-values associated with each coefficient. Significant coefficients are coloured in orange, while non-significant coefficients are shown in green. **(B)** Volcano plot illustrating coefficients computed for DNA methylation data using the genomic integration model. The interpretation follows the same convention as panel (A). **(C)** Dot plot representing enrichment analysis results obtained using significantly regulated genes identified through CNV analysis. The X-axis displays the gene ratio (defined as the number of significant genes within the pathway divided by the cardinality of the pathway) associated with each KEGG pathway, while the Y-axis lists the pathway names. Dot size corresponds to the number of significant genes within each pathway, and color intensity represents the P-value significance. **(D)** Dot plot depicting results from DNA methylation enrichment analysis. The interpretation follows the same convention as panel (C).

Conversely, we found that 38% of the genes have the expression significantly regulated by methylation events, with 68% of these displaying negative coefficients (Figure 2B). This suggests that higher methylation levels generally suppress gene expression, as expected, but that a small proportion of genes have positive coefficients indicating that, although methylated, their expression is still high, that might be explained by the presence of distal unmethylated regulatory elements or of transcription factors resistant to methylation. These genes are found to play a role in immune escape, cell adhesion and neuroactive ligand-receptor interaction (as reported in Figure 2D).

Finally, there are 4230 genes that are significantly regulated by both CNV and methylation simultaneously that modulate gene expression both through a compensatory mechanism (where genes with multiple copies are methylated), or conversely, by enhancing each other’s impact (where genes with multiple copies are unmethylated).

Among them we found SCYL3 (SCY1-Like Pseudokinase 3), FGR (FGR Proto-Oncogene, Src Family Tyrosine Kinase) and GCLC (Glutamate-Cysteine Ligase Catalytic Subunit). SCYL3 is located on chromosome 1q24.2 and encodes a protein that has a kinase domain and four HEAT repeats. SCYL3 is also known as protein-associating with the carboxyl-terminal domain of ezrin (PACE-1). This protein is known to be overexpressed in cancers of the liver, breast and colon and has already been associated with cancer growth^40^. FGR has already been associated with Ovarian Cancer, since it accelerates peritoneal seeding, and potentiates an immunosuppressive immune landscape that facilitates ovarian cancer progression and metastasis^41^. GCLC is the catalytic subunit of the GCL enzyme, containing the active site responsible for the bond formation between the amino group of cysteine and the γ-carboxyl group of glutamic acid needed for glutathione (GSH) synthesis. Several reports already associated high GSH levels with cisplatin or carboplatin resistance in Ovarian Cancer through drug uptake reduction and increased intracellular drug detoxification/ inactivation, increased DNA repair and inhibition of apoptosis drug-induced oxidative stress^42^.

To visually summarize the global integration, in Figure 3 the circos plot provided by the shinyApp, gives a complete overview of the interplay between gene expression, CNV and methylation levels. We identified most of the altered chromosomal regions known to characterise high grade ovarian cancer such as 8q24 (with MYC and PVT1 genes), 3q26 (with PIK3CA and TERC genes), 19q12 (with CCNE1) and 5p15 (with TERT) for gains and 17p13 (with TP53 gene), 13q14 (with RB1) and 6q26 (with ARID1B gene) for losses (Figure 3A-C).

**Figure 3.**
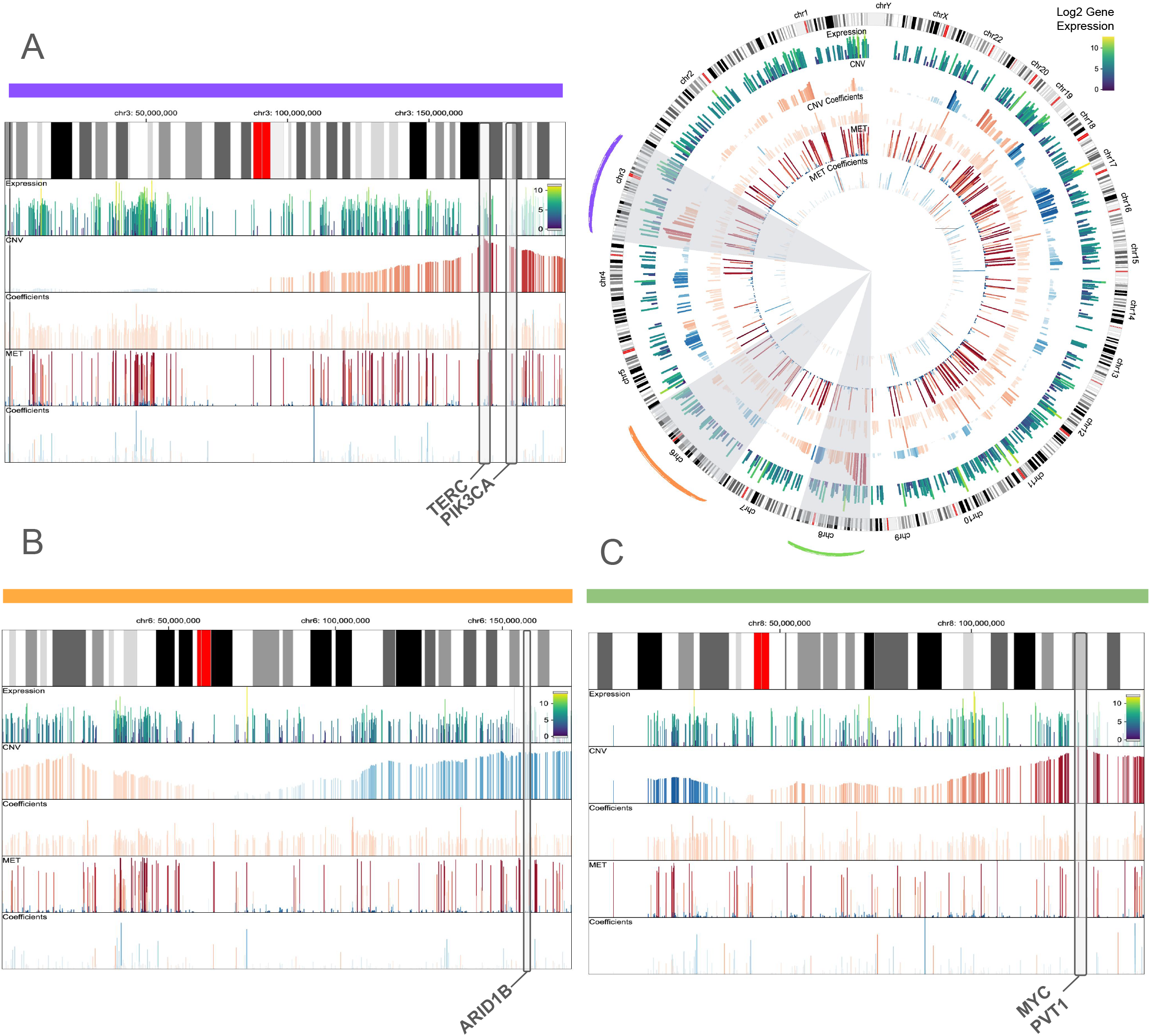
Comprehensive integration analysis of the TCGA OV dataset. The Circos plot provides an overview of the genomic integration models. Along each chromosome, the inner layers display: gene expression values (first layer), copy number profiles (second layer), coefficient estimates from the expression–CNV model (third layer), DNA methylation beta values (fourth layer), and coefficient estimates from the expression– methylation model (fifth layer). **(A)** Linear view of chromosome 3, highlighting the positions of TERC and PIK3CA. **(B)** Linear view of chromosome 6, highlighting the position of ARID1B. **(C)** Linear view of chromosome 8, highlighting the positions of MYC and PVT1.

In particular, most genes located on the q24 arm of chromosome 8 are highly expressed (Figure 3C). However, some of them, despite having high copy numbers, exhibit significant methylation, such as PVT1, which leads to a reduction in its expression. On chromosome 3 (Figure 3A), the q26 arm reveals a large number of highly expressed genes, driven by gain events like PIK3CA. However, some genes, such as TERC, are also modulated by methylation. In contrast, chromosome 6 (Figure 3B) contains a large region of losses, where we observe AIRD1B, whose expression is regulated not only by copy number variation (CNV) but also by methylation levels.

After accounting for the alterations due to CNV and methylation, we evaluated the transcriptional activity of TF, conditioned on the genomic alterations.

The three transcription factors that appear to be most active in ovarian cancer are TAL1, FOXP3, and GATA1 (Figure 4A-B). TAL1, located on chromosome 1 arm q33, is known to regulate cell differentiation and proliferation. While its precise role in ovarian cancer is still under investigation, evidence suggests that TAL1 contributes to resistance to apoptosis ^43^ and to the evasion of immune surveillance. Under physiological conditions, TAL1 functions exclusively in the nucleus where it forms transcriptional complexes with other factors. Our results indicate an enrichment of target genes localised in the cytoplasm and in the membrane (Figure 4D), suggesting an aberrant localization of TAL1 in ovarian cancer cells that might contribute to an extensive disruption of its regulatory program. FOXP3 and GATA1 are both located on the X chromosome (Xq11.23) and their expression is enhanced due to their location in a chromosomal region characterised by gain events. FOXP3 promotes immune evasion by enhancing the activity of Tregs, thereby allowing cancer cells to avoid immune surveillance and contributing to tumor progression^44^. In our results, FOXP3 positively regulates, among others, TGFB1, STAT4, and TLR2 all transcription factors known to play key roles in immune and inflammatory response influencing the tumor microenvironment^45–47^ (Figure 4C). GATA1 on the other hand is known to regulate hematopoiesis and erythropoiesis^48^ and it may influence tumor cell proliferation and survival through the modulation of genes such as MYC (Figure 4E).

**Figure 4.**
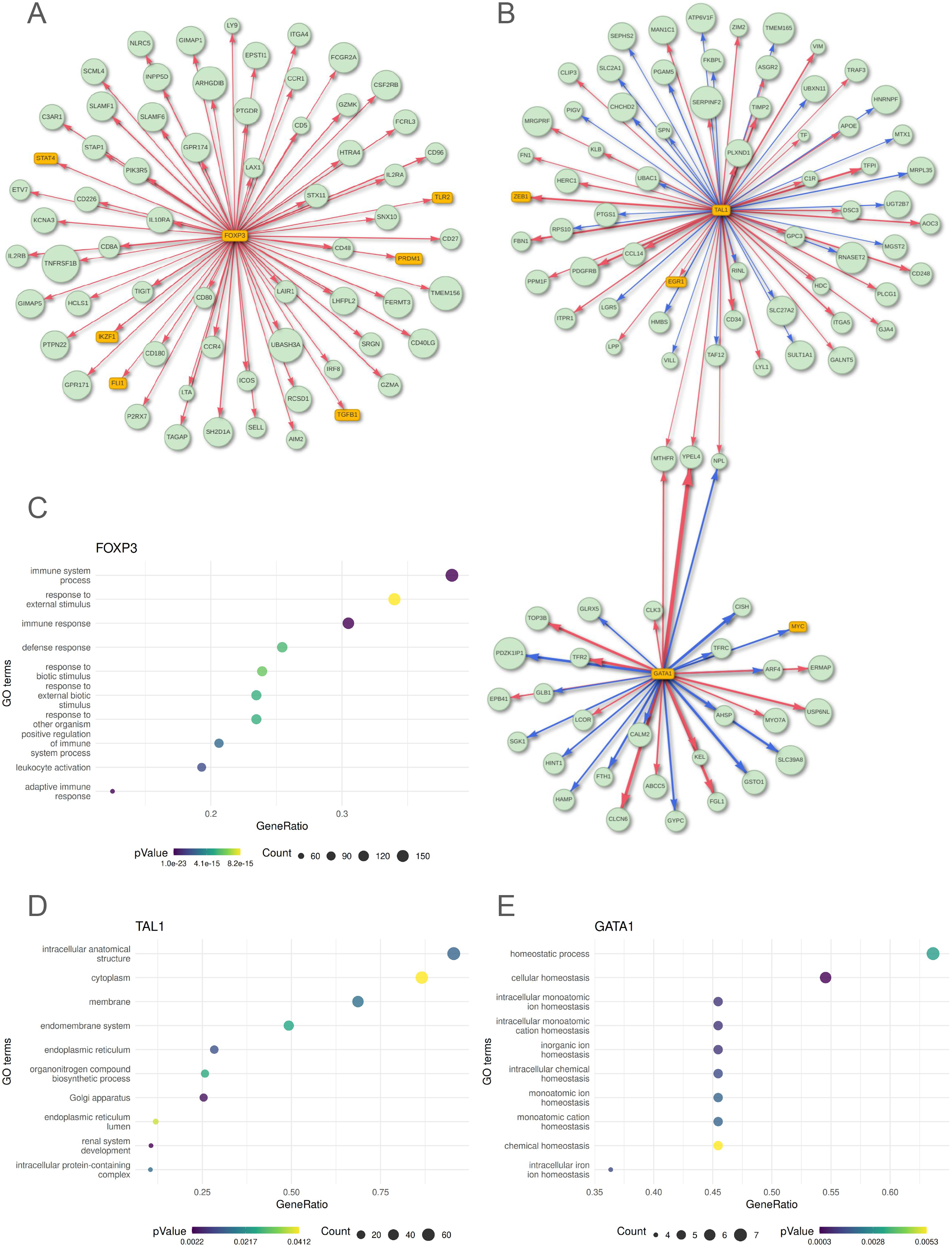
Transcriptional regulation analysis on the TCGA OV dataset. **(A)** Network illustrating significant interactions involving FOXP3, as identified by our transcriptional integration model. Yellow rectangles represent transcription factors (TFs), green circles indicate target genes, and edges denote significant regulatory interactions. Red arrows indicate positive regulation, while blue arrows represent repression. **(B)** Network showing significant interactions for TAL1 and GATA1, following the same graphical conventions as in panel (A). **(C–E)** Dot plots showing Gene Ontology (GO) enrichment analysis of the target genes regulated by FOXP3 (C), TAL1 (D), and GATA1 (E). The x-axis indicates the gene ratio (i.e., the number of significant genes within each GO term divided by the total number of genes in that term), while the y-axis lists the GO term names. Dot size reflects the number of significant genes, and color intensity represents the significance of the enrichment (p-value).

### gINTomics browses through the prognostic transcriptional rewiring comparing patients with long and short-term survival

Ovarian cancer is known for its high aggressiveness and poor prognosis, largely due to its tendency to be diagnosed at an advanced stage. The prognosis varies significantly among patients and comparing molecular profiles of long-surviving and short-surviving patients is crucial for identifying biomarkers and mechanisms that influence disease progression. In this case we selected samples from the TCGA-OV cohort and divided them in two groups according to their Progression Free Survival time (PFS), PFS shorter than 6 months (n=29), and PFS longer than 5 years (n=21).

Globally, we found a total of 1064 DEGs (FDR<=0.1), of which 202 are significantly deregulated by genomic alterations (64 by CNV, 100 by methylation and 38 by both). In general we found many tumour suppressors significantly downregulated in short-term survivors by deletion or methylation events. Among them we were able to identify genes that are known to play a key role in cancer, such as PAX2^49^, FRZB^50^, CDO1^51^, LRIG1^52^, KL^53^ and others. It is interesting to note how PAX2 (Figure 5A) is characterised by a higher CNV in long compared to short survival that reflects in a higher expression of PAX in patients with a better prognosis.

**Figure 5.**
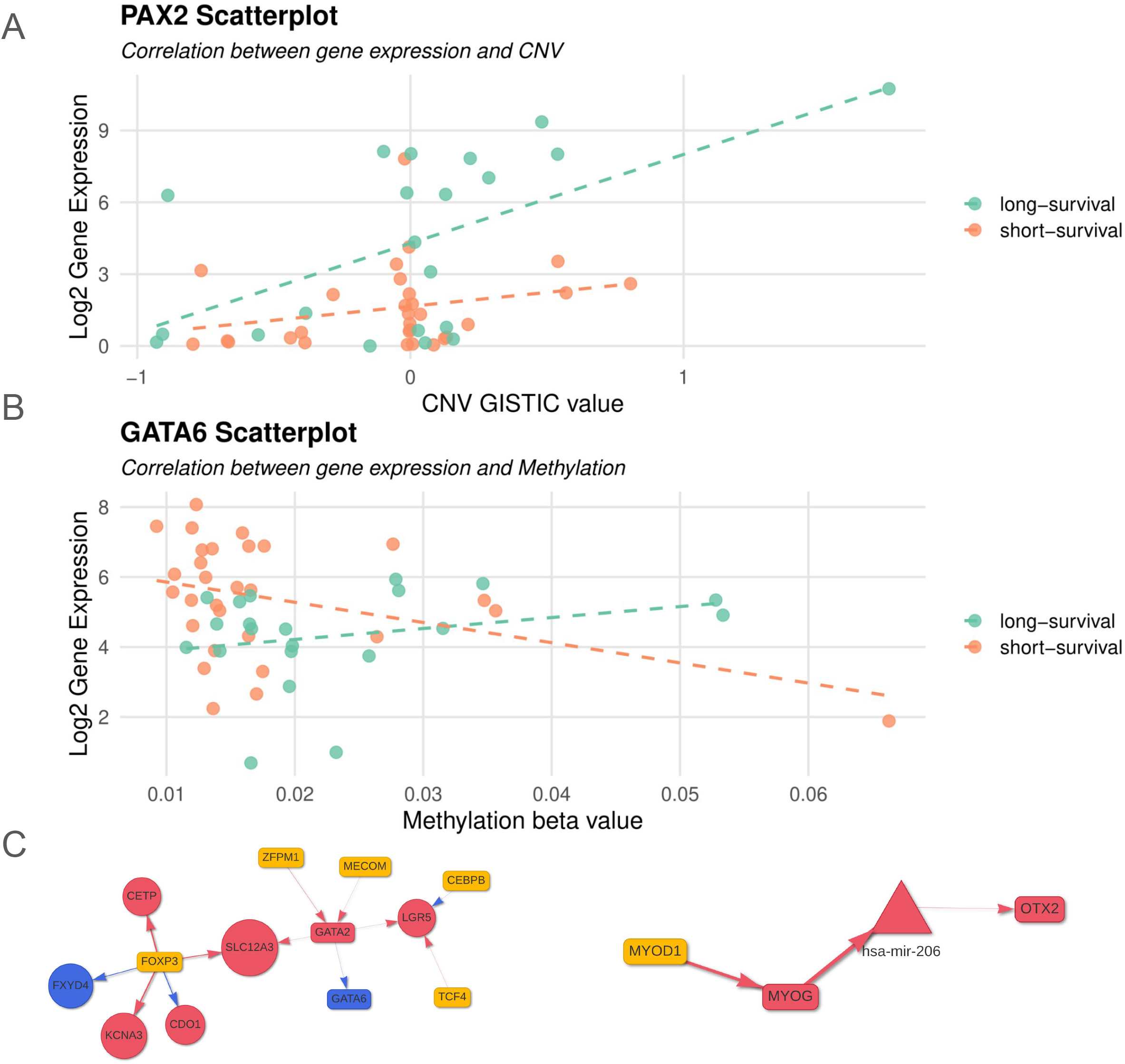
Integrative analysis comparing short- and long-term survival in the TCGA OV dataset. **(A)** Scatter plot showing the relationship between CNV and gene expression for the PAX2 gene in long-term (green) and short-term (orange) survival groups. The x-axis represents CNV GISTIC values, while the y-axis shows log2-transformed gene expression. A regression line is fitted for each group. **(B)** Scatter plot showing the relationship between DNA methylation and gene expression for the GATA6 gene in long-term (green) and short-term (orange) survival groups. The interpretation follows the same convention as panel (A), except that the x-axis represents methylation beta values. **(C)** Network illustrating significant interactions between transcription factors and differentially expressed genes (DEGs) between long- and short-term survival groups. Rectangles represent TFs, circles represent target genes. Red nodes indicate genes downregulated in the short-term survival group, while blue nodes indicate upregulated genes. Red arrows denote positive regulation; blue arrows denote repression.

On the other hand we found different overexpressed ovarian cancer oncogenes that are located in amplification or hypomethylated regions and significantly regulated by their genomic status, such as FOXO1^54^, GATA6^55^, FXYD4^56^ and others. In particular GATA6 (Figure 5B) is down-regulated in long survival and shows a differential association with its methylation status in the two classes.

Looking at the differential CNV we detected 412 differentially altered genes (P Value <= 0.05), while the differential Methylation analysis identified 71 differentially methylated genes whose methylation significantly impacted gene expression in the short-term survival group.

Moreover our transcriptional analysis (Figure 5C) revealed an interesting interaction between GATA2 and GATA6. In particular, GATA2, which is downregulated in the short-term survival group, negatively regulates the expression of GATA6, which is in fact upregulated in the same group. Interestingly, high expression levels of GATA6 have already been associated with worse prognosis in Ovarian Cancer, while high expression of GATA2 with better prognosis^57^. Among the most interesting biomarkers found to be upregulated in long-term survival group, we have hsa-miR-206 (logFC = 5, p.adjust = 0.007), this miRNA has been previously found to be down-regulated in epithelial ovarian cancer tissues and cell lines. In particular, hsa-miR-206 inhibits the development of epithelial ovarian cancer by directly targeting c-Met and inhibiting the c-Met/AKT/mTOR signaling pathway^58^. Interestingly, our analysis revealed that this miRNA is positively regulated by MYOG and its expression is also positively related to OTX2 expression; all these genes are upregulated in the long-term survival group.

Finally, for the short-term survival group, we found other interesting biomarkers such as hsa-miR-34b (logFC = −2.5, p.adjust = 0.07); FXYD4 (logFC = −3.2, p.adjust = 0.027) which is strongly regulated by both CNV and methylation; TRIM15 (logFC = −2.3, p.adjust = 0.048); TRIM9 (logFC = −2.2, p.adjust = 0.056) and others.

Taken together all these results highlight the robustness of `gINTomics` in the identification of the transcriptional and genomic alterations responsible for Ovarian Cancer prognosis. The presence in our results of genes that have already been associated with Ovarian Cancer prognosis further support the reliability of our models.

## Conclusions

`gINTomics` represents a significant advancement in multi-omics data integration by providing a hierarchical framework that captures the directional flow of biological information across molecular layers. A key strength of the package lies in its flexibility to accommodate different combinations of omics data types, from simple two-omics integrations (such as CNV-expression or methylation-expression) to comprehensive multi-omics analyses incorporating genomics, transcriptomics, epigenetics, and miRNA data. This adaptability allows researchers to work with diverse experimental settings and data availability. Validation on TCGA ovarian cancer data demonstrated the framework’s ability to identify biologically relevant regulatory mechanisms and prognostic biomarkers, confirming the effectiveness of the multi-staged approach. The integration of an interactive Shiny application with advanced visualization tools, including circos plots, makes `gINTomics` accessible to both bioinformaticians and researchers without coding skills. The framework’s capability to simultaneously handle genomic, epigenetic, and transcriptional alterations while identifying complex regulatory networks positions `gINTomics` as a valuable tool for integrative analyses and biomarker discovery. Its implementation as a Bioconductor package ensures reproducibility and accessibility to the scientific community.

---

**Supplementary Table 1**.

A selection of the most widely used multi-omic integration methods that are available within the R and Bioconductor ecosystem. For each method, the name, analytical approach, and primary objective are reported. The list is not exhaustive, but highlights the most commonly adopted tools in this computational environment. A comprehensive list of packages for multi-omics integration is also available here https://github.com/mikelove/awesome-multi-omics.

**Table 2.** Overview of available functions for omics data integration. This table summarizes the `gINTomic` functions applicable to the omics data types, highlighting the available approaches for multi-omics integration.

| **gINTomic**<br>**Integration functions** | Gene<br>Expression Data | miRNA<br>Expression Data | Copy Number<br>Variation Data | Methylation<br>Data |
| --- | --- | --- | --- | --- |
| run_cnv_integration | ✓ |  | ✓ |  |
| run_cnv_integration |  | ✓ | ✓ |  |
| run_met_integration | ✓ |  |  | ✓ |
| run_met_integration |  | ✓ |  | ✓ |
| run_genomic_integration | ✓ |  | ✓ | ✓ |
| run_tf_integration | ✓ | ✓ |  |  |
| run_tf_integration | ✓ |  |  |  |
| run_tf_integration | ✓ | ✓ |  |  |
| run_multiomics | ✓ | ✓ | ✓ | ✓ |

