## Supplemental Table 1 for "From Genomics Alterations to Expression Dynamics: A Hierarchical Multi-Omics Integration Framework with gINTomics"

| **Method** | **Meta-dimensional** | **Outcome** | **Availability** |
| --- | --- | --- | --- |
| **MOFA^10^** | inferred latent factors | Clustering, dimensionality reduction | BioC |
| **Mixomics^11^** | multivariate dimension reduction | Clustering, dimensionality reduction, signature evaluation | BioC |
| **MOVICS^12^** | multi-omics clustering algorithms | Clustering | BioC |
| **MOFA+^13^** | extension of MOFA for single cell data | Clustering, dimensionality reduction, inference of differentiation trajectories | BioC |
| **MOSClip^14^** | topological pathway analysis | Survival-associated gene modules. | BioC |
| **MOSS^15^** | sparse singular value decomposition | Clustering, non-linear embedding | GitHub |
| **Online iNMF^16^** | online integrative non-negative matrix factorization | dataset alignment and clustering | CRAN, GitHub |
| **MIMaL^17^** | tree-based regression model (SHAP) | Predict metabolite changes from proteomic changes | <https://mimal.app> |
| **MOMA^18^** | VIPER, DIGGIT Analysis, Bayesian Integration, Saturation Analysis, Modularity Analysis | Identification of Master Regulator Blocks and sample stratification | BioC |
| **OMICsPCA^20^** | Principal Component Analysis | analysis of overall distribution of OMICs assays, grouping assays, identification of source of variation | BioC |
| **timeOmics^21^** | linear mixed model, unsupervised clustering | identification of molecular features strongly associated with time | BioC |
| **MoNETA^22^** | Multi-omics Network Embedding | Sample sub-typing, cell types and functional states characterization | GitHub |
| **Method** | **Hierarchical methods** | **Outcome** |  |
| **COSMOS^19^** | Integer Linear Programming optimization strategy | Estimate the activities of transcription factors and kinases, network-level causal reasoning | BioC |
| **Zhu et al. 2016^23^** | Penalized regression | Biomarker associated to clinical outcome | **-** |
| **Wu et al. 2019^24^** | Penalized regression | Biomarker associated to clinical outcome | **-** |
| **iBAG^25^** | integrative Bayesian model | infer biological relationships among omics and association with clinical outcome | - |

**Supplementary Table 1. Selected multi-omic integration methods available in R and Bioconductor**
A selection of the most widely used multi-omic integration methods that are available within the R and Bioconductor ecosystem. For each method, the name, analytical approach, and primary objective are reported. The list is not exhaustive, but highlights the most commonly adopted tools in this computational environment. A comprehensive list of packages for multi-omics integration is also available here <https://github.com/mikelove/awesome-multi-omics>
